# Colonic Oxygen Microbubbles Augment Systemic Oxygenation and CO_2_ Removal in a Porcine Smoke Inhalation Model of Severe Hypoxia

**DOI:** 10.1101/2021.12.08.466665

**Authors:** Paul. A. Mountford, Premila. D. Leiphrakpam, Hannah. R. Weber, Andrea McCain, Robert. M. Scribner, Robert. T. Scribner, Ernesto M. Duarte, Jie Chen, Mark. A. Borden, Keely. L. Buesing

**Affiliations:** Respirogen, Inc., Boulder, Colorado, USA; University of Nebraska Medical Center, Omaha, Nebraska, USA; University of Florida College of Medicine, Gainesville, Florida, USA; University of Colorado Boulder, Boulder, Colorado, USA

## Abstract

Inhalation injury can lead to pulmonary complications resulting in the development of respiratory distress and severe hypoxia. Respiratory distress is one of the major causes of death in critically ill patients with a reported mortality rate of up to 45%. The present study focuses on the effect of oxygen microbubble (OMB) infusion via the colon in a porcine model of smoke inhalation-induced lung injury. Juvenile female Duroc pigs (n=6 colonic OMB, n=6 no treatment) ranging from 39-51 kg in weight were exposed to smoke under general anesthesia for 2 h. Animals developed severe hypoxia 48 h after smoke inhalation as reflected by reduction in SpO2 to 66.3 % ± 13.1% and P_a_O_2_ to 45.3 ± 7.6 mmHg, as well as bilateral diffuse infiltrates demonstrated on chest x-ray. Colonic OMB infusion (75 – 100 mL/kg dose) resulted in significant improvements in systemic oxygenation as demonstrated by an increase in P_a_O_2_ of 13.2 ± 4.7 mmHg and SpO_2_ of 15.2% ± 10.0% out to 2.5 h, compared to no-treatment control animals that experienced a decline in P_a_O_2_ of 8.2 ± 7.9 mmHg and SpO_2_ of 12.9% ± 18.7% over the same timeframe. Likewise, colonic OMB decreased P_a_CO_2_ and P_mv_CO_2_ by 19.7 ± 7.6 mmHg and 7.6 ± 6.7 mmHg, respectively, compared to controls that experienced increases in P_a_CO_2_ and P_mv_CO_2_ of 17.9 ± 11.7 mmHg and 18.3 ± 11.2 mmHg. We conclude that colonic OMB therapy has potential to treat patients experiencing severe hypoxemic respiratory failure.

**One Sentence Summary:** Enteral oxygen microbubbles increase systemic oxygen and decrease carbon dioxide levels in acutely hypoxic pigs after smoke inhalation-induced respiratory failure.

## Introduction

Prior to the spread of SARS-CoV-2, acute respiratory distress syndrome (ARDS) historically occurred in ∼10% of patients entering the intensive care unit – affecting nearly 190,000 patients per year in the US alone – with a reported mortality rate ranging from 35% to 46% *(1, 2)*. As of October 20, 2021, a total of 730,368 COVID-19 deaths have been reported in the US *(3)*. The primary symptom of COVID-19 infection requiring hospitalization is hypoxemic respiratory failure*(4)*. Regardless of underlying pathology, mechanical ventilation remains the mainstay of oxygenation and ventilatory support for severe respiratory failure; however, complications such as ventilator-induced lung injury, ventilator-associated pneumonia, barotrauma, and progressive deconditioning leading to ventilator dependence remain unacceptably high *(5)*. Despite decades of research, therapeutic options for patients with severe hypoxemia that fail mechanical ventilatory support are limited.

Extracorporeal membrane oxygenation (ECMO) – a temporary, artificial extracorporeal support of the respiratory and/or cardiac system – is one of the last resorts for treating refractory respiratory failure *(6)*. On the spectrum of ARDS treatment, ECMO is invasive and complex. When used for respiratory failure, it serves as a pulmonary bypass, circulating the patient’s blood through an external circuit to exchange oxygen and carbon dioxide *(7)*. While usually effective at correcting acute hypoxemia, ECMO has a substantial contraindication list and risk profile leading to unacceptably high 30- and 60-day mortality rates of 39-50% *(6, 8)*. The risks of ECMO are significant, and include hemorrhage, thrombocytopenia, circuit failure, embolism, hemolysis, and limb ischemia among other potential complications *(8)*. Alternative therapies that can provide meaningful systemic oxygen augmentation and allow the lungs to rest without the risks of ECMO need to be actively pursued and investigated.

Oxygen microbubble (OMB) therapy is a novel technology that shows promise as a method of extrapulmonary oxygenation that is relatively simple and safe to administer, does not require the use of anticoagulants and does not have the risk profile associated with ECMO. OMB is comprised of a high concentration of micron-scale (1-20 um diameter) bubbles that contain an oxygen “core” and are encapsulated by a lipid monolayer shell *(9)*, similar to the pulmonary alveolus. When administered as a bolus dose into the abdominal cavity (akin to peritoneal dialysis), OMBs have been reported to augment systemic oxygenation and improve outcome in small animal pilot studies involving unilateral pneumothorax, tracheal occlusion and LPS-mediated severe ARDS *(9–11)*.

Here, we introduce a novel delivery pathway for OMB therapy - the colon - as an improved translational candidate for minimally invasive, nonsurgical oxygenation and carbon dioxide removal for the treatment of severe hypoxia. The colonic mucosa is associated with a rich capillary matrix. Oxygen tension in the mucosal layer has been studied in small animal models of hyperbaric oxygen therapy, where investigators found that oxygen diffused from intestinal tissue and established a radial gradient from the tissue interface to the colonic lumen *(12)*. It naturally follows that if systemic hyperoxia can augment luminal oxygen content via the capillary gradient, establishing an elevated oxygen content in the colonic lumen would lead to diffusion across the capillary bed, augmenting systemic oxygenation in states of hypoxemia. A similar argument holds for carbon dioxide exhaust from the same capillary gradient. Moreover, the colon provides an ability to deliver a clinically relevant volume of OMB without the need to place a surgical port, as required by alternative enteral routes. This study examines the hypothesis that colonic OMB therapy can significantly increase systemic oxygen levels, and reduce systemic carbon dioxide levels, in a large-animal model of severe hypoxia.

## Results

### Lung injury due to smoke inhalation injury

Prior to OMB treatment at 48 h after smoke inhalation, lung injury was assessed by chest x-ray (CXR) and carotid, femoral and pulmonary arterial catheter blood gas sampling. CXR confirmed the presence of diffuse bilateral infiltrates indicative of ARDS (Fig. 1A,B,D and E). While on a fraction of inspired oxygen (FiO2) level of 21%, P_a_O_2_ decreased from 93.8 ± 26.5 mmHg to 45.3 ± 7.6 mmHg and SpO_2_ dropped from 97.0% ± 2.1% to 66.3% ± 13.1%, while P_a_CO_2_ rose from 42.0 ± 4.3 mmHg to 58.2 ± 4.1 mmHg at 30 min prior to OMB treatment (Fig. 1C). There was an increase in IL-6 inflammation within the lungs (Fig. 1H, BAL). Additionally, we observed a significant increase in overall lung injury score (Fig. 1I) and average wet-dry (W/D) weight ratio of lung tissues 48 h after smoke exposure compared to the control animals (Fig. 1J).

**Fig. 1.**
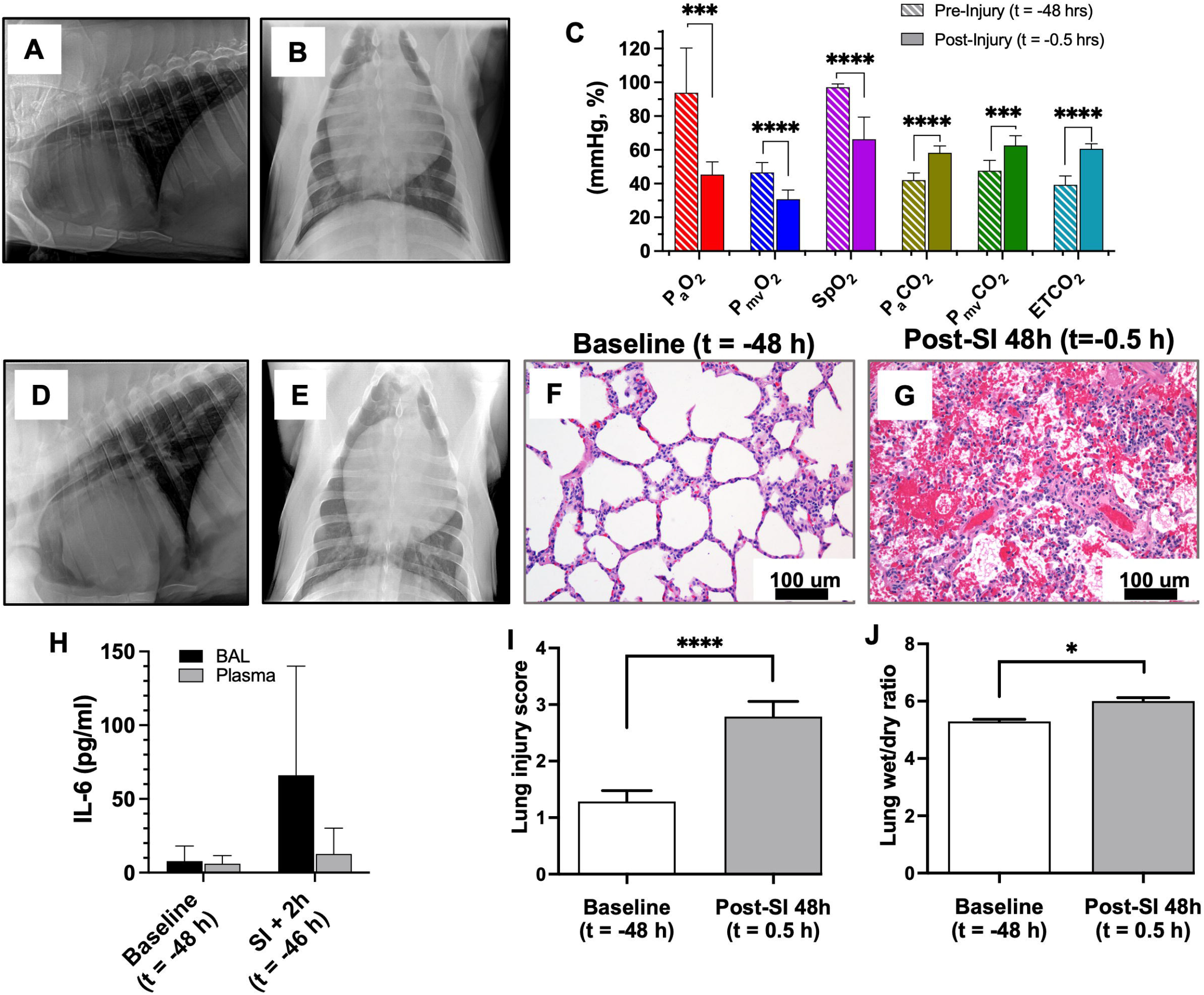
Porcine Smoke Inhalation Injury. **A** and **B**) Before and after (**D** and **E**) chest x-ray images confirming the presence of diffuse bilateral infiltrates indicative of ARDS due to smoke inhalation injury. **C**) P_a_O_2_ (red, *p=*0.000147), P_mv_O_2_ (blue, *p<*0.0001), SpO_2_ (violet, *p<*0.0001), P_a_CO_2_ (gold, *p<*0.0001), P_mv_CO_2_ (green, *p=*0.000123) and ETCO_2_ (teal, *p<*0.0001) measurements taken both before (t=-48 h) and after (t=-0.5 h) smoke inhalation injury. Hematoxylin and eosin (H&E) staining of paraffin embedded lung tissue sections of baseline (t = -48 h) (**F**) and SI + 48 h (t = -0.5 h) (**G**) animals (scale bar = 100 um). **H**) IL-6 marker analysis for baseline and smoke injury (SI) + 2 h (t = -46 h) for BAL and Plasma samples. Comparison of lung injury score (**I**) and lung wet/dry ratios (**J**) showing a significant difference between control and SI + 48 h (t = -0.5 h) samples (*p<*0.0001 and *p=*0.0188, respectively)

### Systemic O_2_ delivery and CO_2_ removal

Upon achieving severe hypoxia due to smoke inhalation injury, OMB was administered to the colon (Fig. 2D) in the form of three bolus injections at a rate of 500 mL/min for a total dose size of 75 to 100 mL/kg (3.6-4.5 L total for pigs ranging from 40-50 kg). The OMB had a number-weighted average microbubble diameter of 1-10 um with most of the oxygen gas volume existing in bubbles 1-20 um in diameter (Fig. 2B). Within 120 min after the start of OMB treatment, all blood and non-invasive oxygen vitals showed statistically higher oxygen content for animals receiving OMB treatment compared to no-treatment control animals. Specifically, P_a_O_2_ rose significantly for OMB-treated animals within the first 15 min to 52.5 ± 6.9 mmHg (OMB Δ P_a_O_2_ = 9.3 ± 6.4 mmHg, no treatment (NT) Δ P_a_O_2_ = -2.3 ± 6.7 mmHg) and continued rising to 56.4 ± 7.9 mmHg (OMB Δ P_a_O_2_ = 13.2 ± 4.7 mmHg, NT Δ P_a_O_2_ = -8.2 ± 7.9 mmHg) after 150 min (Fig. 3A,C). Additionally, P_mv_O_2_ increased significantly for OMB treatment animals to 34.8 ± 5.8 mmHg (5.0 ± 5.9 mmHg over -7.3 ± 6.9 mmHg as seen by the NT animals) after 150 min (Fig. 3A,D). SpO_2_ also rose significantly by 14.2% ± 9.5% after 60 min as compared to the drop seen in the no-treatment group (NT ΔSpO_2_ = -4.2% ± 14.91%) and was statistically higher at 150 min (OMB ΔSpO_2_ = 15.2 ± 10.0 %, NT ΔSpO_2_ = -12.9 ± 18.7 %, Fig. 3A,E).

**Fig. 2.**
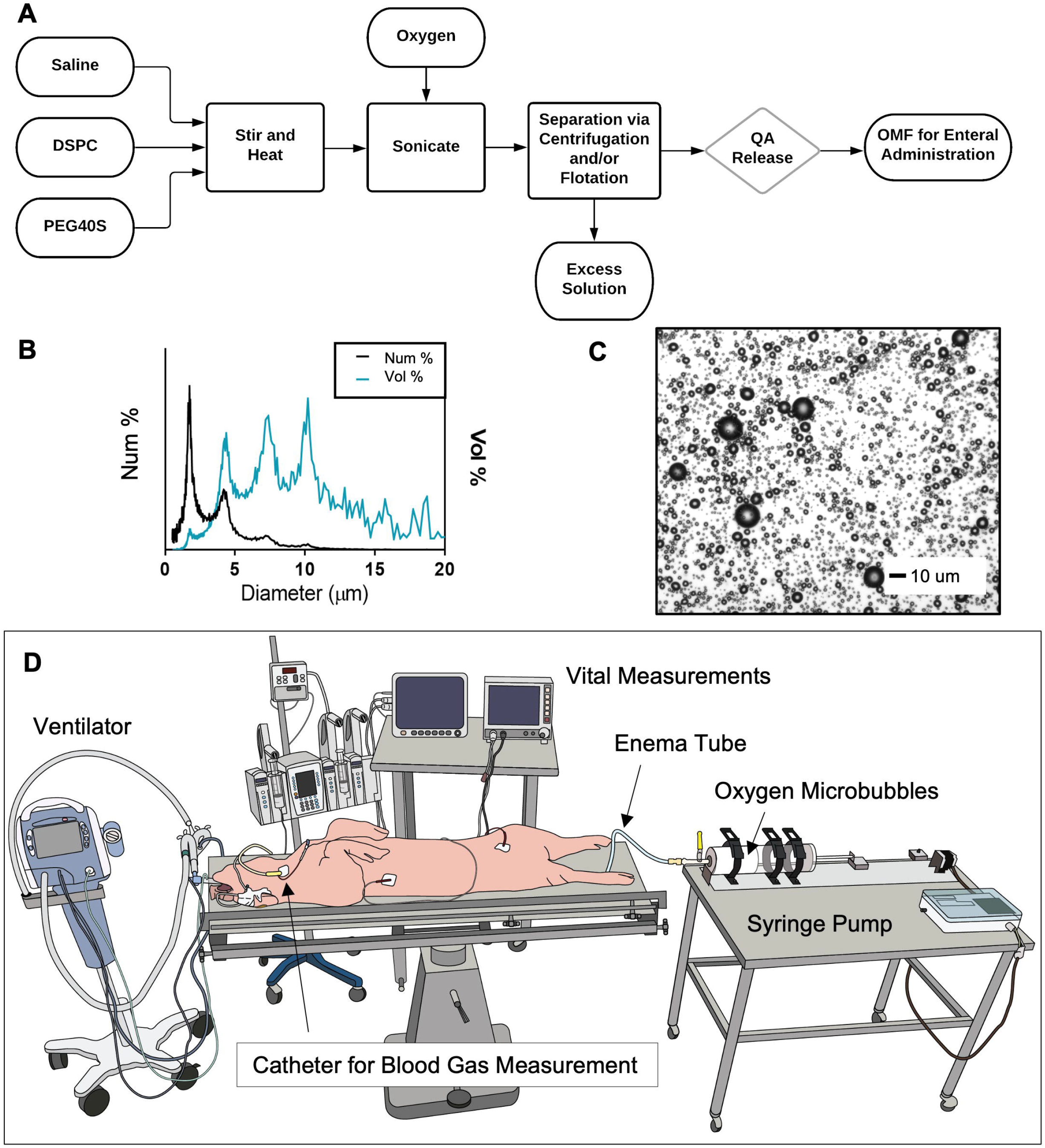
Colonic OMB Administration. **A**) Process flow diagram showing the production process for creating OMB via sonication and differential centrifugation. **B**) Particle size by both number percent (black) and volume percent (blue) frequency for the OMB samples. **C**) Microscopy image showing the size of the OMB (scale bar = 10 um). **D**) Schematic showing the colonic delivery of OMB to a smoke inhalation lung injured pig on minimal mechanical ventilation (FiO_2_=0.21).

**Fig. 3.**
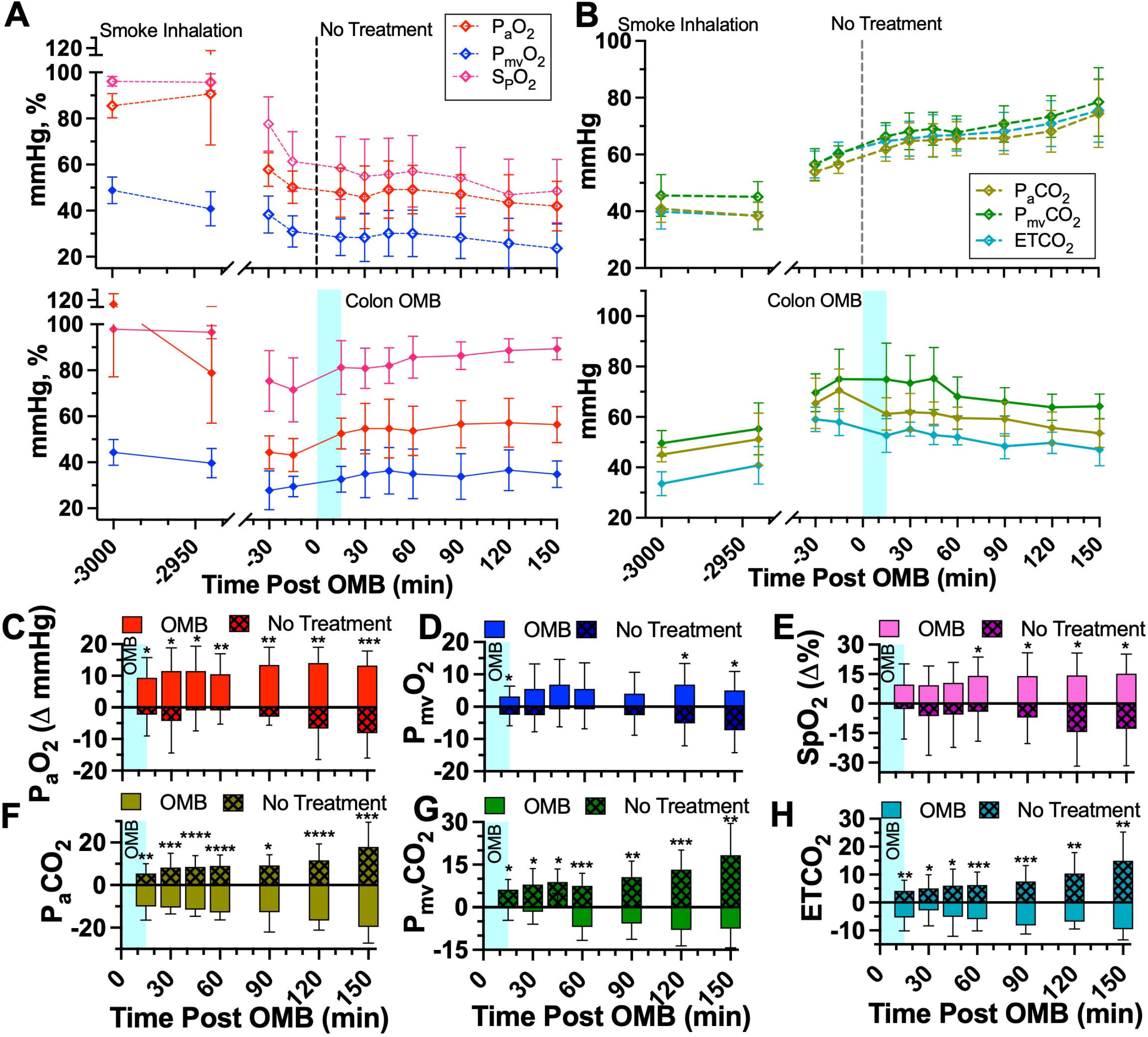
Colonic OMB Systemic Oxygenation and CO2 Removal. **A)** Blood oxygen and **B**) CO_2_ measurements for NT (top) and colonic OMB (bottom) treatment groups from before smoke inhalation injury (t = -3000 min) out to t = 150 min post treatment time. The change in oxygen vitals for NT (hashed) and OMB (solid) treatment groups showing the statistical significance at times t = 15, 30, 45, 60, 90, 120 and 150 min ((**C**) P_a_O_2_ (red), (*p=*0.0117, 0.0124, 0.0137, 0.0058, 0.0014, 0.0022 and 0.0005), (**D**) P_mv_O_2_ (blue), (*p=*0.0138, 0.0597, 0.0799, 0.1568, 0.1230, 0.0171 and 0.0110), (**E**) SpO_2_ (violet), (*p=*0.1350, 0.1239, 0.0788, 0.0329, 0.0212, 0.0095 and 0.0133), (**F**) P_a_CO_2_ (gold), (*p=*0.0025, 0.0004, <0.0001, <0.0001, 0.0109, <0.0001 and 0.0003), (**G**) P_mv_CO_2_ (green), (*p=*0.0330, 0.0113, 0.0132, 0.0007, 0.0030, 0.0009 and 0.0018) and (**H**) ETCO_2_ (teal), (*p=*0.0040, 0.0269, 0.0136, 0.0007, 0.0004, 0.0015 and 0.0012)).

Moreover, arterial measurements showed blood gas CO_2_ declining from treatment until the end of the study for animals receiving OMB. Specifically, P_a_CO_2_ (Fig. 3B,F) was significantly lower for animals receiving OMB after 15 min at 61.20 ± 6.39 mmHg (OMB Δ P_a_CO_2_ = -10.0 ± 6.4 mmHg, NT Δ P_a_CO_2_ = 5.4 ± 4.6 mmHg) and remained lower out to 150 min at 53.6 ± 5.6 mmHg (OMB Δ P_a_CO_2_ = -19.7 ± 7.6 mmHg, NT Δ P_a_CO_2_ = 17.9 ± 11.7 mmHg). P_mv_CO_2_ (Fig. 3B,G) was statistically lower for animals receiving OMB therapy at 15 min (OMB Δ P_mv_CO_2_ = -0.2 ± 4.4 mmHg, NT Δ P_mv_CO_2_ = 6.2 ± 3.5 mmHg) and 150 min (OMB Δ P_mv_CO_2_= -7.6 ± 6.7 mmHg, NT Δ P_mv_CO_2_ = 18.3 ± 11.2 mmHg). The ETCO_2_ measurements showed a similar decline for the animals receiving OMB from 15 min (OMB Δ ETCO_2_ = -5.3 ± 4.8 mmHg, NT Δ ETCO_2_ = 4.2 ± 3.8 mmHg) out to 150 min (OMB Δ ETCO_2_ = -9.6 ± 3.9 mmHg, NT Δ ETCO_2_ = 15.0 ± 10.2 mmHg, Fig. 3B,H).

### Effect of OMB treatment on lung parenchyma, inflammatory cytokines and proteomics

No significant changes were observed in lung injury score and wet/dry weight ratio at the 3-h time point (Fig. 4A-D). However, we observed a significant increase in IL-1β levels in immunoblotting of lung tissue lysates at 48 h post smoke inhalation in smoke injury (SI) animals compared to the control animals (Fig. 4E,F). Expression status of both cytokines was reversed at 3 h post OMB treatment (Fig. 4F), indicating an anti-inflammatory effect of OMB. There were lower IL-6 levels for the OMB group (BAL = 1.7 ± 1.5 pg/mL, Plasma = 5.3 ± 5.5 pg/mL) as compared to the NT group (BAL = 32.9 ± 24.0 pg/mL, Plasma = 14.0 ± 13.9 pg/mL) for both the BAL and Plasma samples (Fig. 4G).

**Fig. 4.**
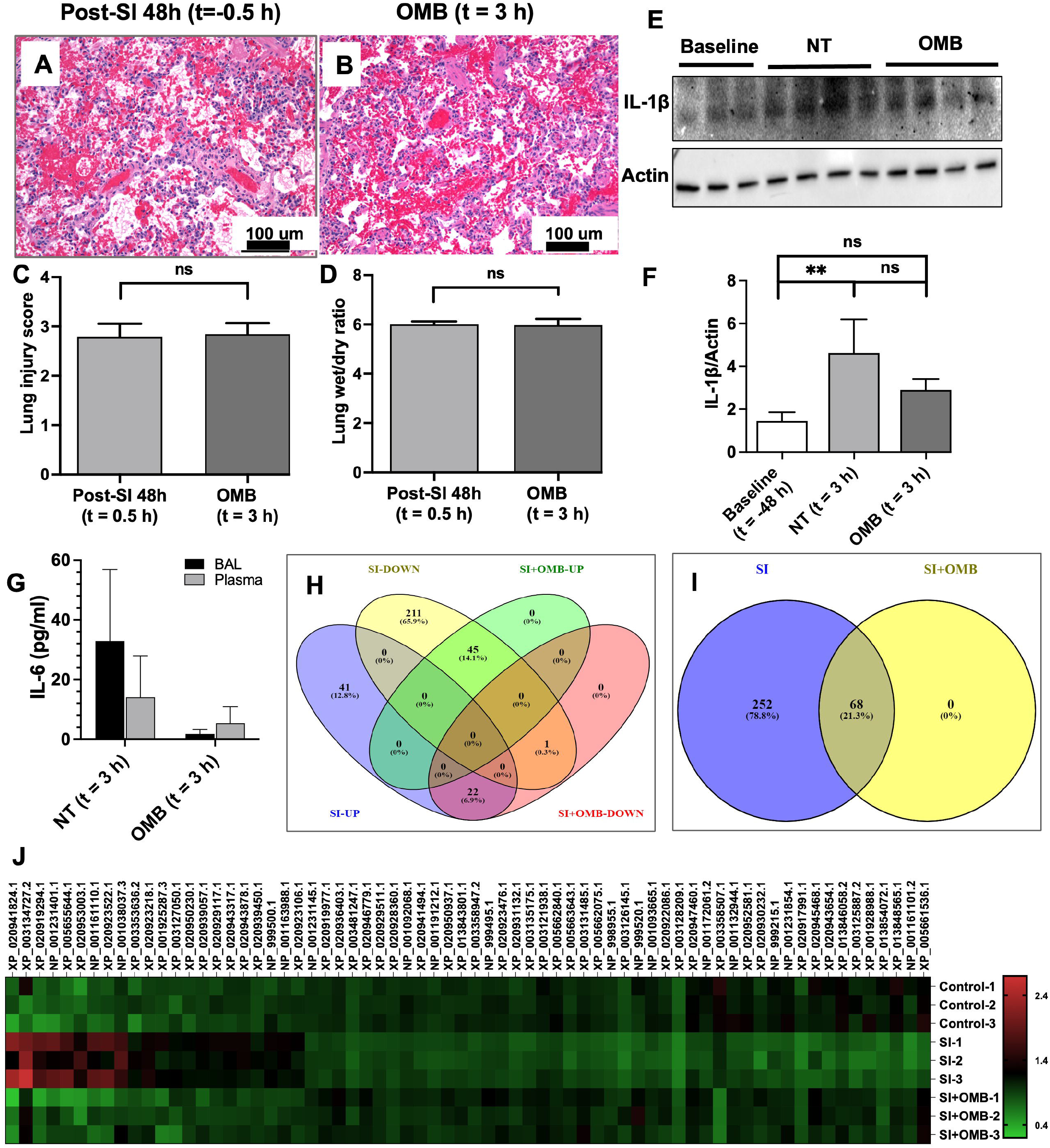
Local and Systemic Injury and Inflammation. Hematoxylin and eosin (H&E) staining of paraffin embedded lung tissue sections of SI + 48 h (t = -0.5 h) (**A**) and OMB treatment (t = 3 h) (**B**) animals (scale bar = 100 um). Comparison of lung injury score (**C**) and lung wet/dry ratios (**D**) between SI + 48 h (t = -0.5 h) and OMB treatment (t = 3 h) samples. **E** and **F**) Immunoblot analysis of IL-1β expression levels in fresh frozen lung tissues of baseline (t = -48 h), NT and OMB samples (t = 3 h). Difference between baseline and NT groups was statistically significant (*p*=0.0092). G) IL-6 marker analysis for NT and OMB (t = 3 h) for BAL and Plasma samples. **H, I and J**) Proteomic analysis of control (t = -48 h), SI (NT, t = 3 h) and SI+OMB (t = 3 h) groups for their global protein expression as described in Material and Methods section. Venn diagram showed 320 proteins with significant differential expression between SI and control groups (**H** and **I**). Heat map analysis of 68 proteins that were significantly upregulated or downregulated at 3 hours post OMB treatment (**J**). A p value of < 0.05 was considered statistically significant.

Fresh frozen lung tissues of control, SI and SI with OMB groups were compared for their global protein expression by proteomic analysis. Several proteins were differentially expressed between these groups. Of these proteins, 320 proteins were significantly upregulated or downregulated at 48 h post smoke exposure compared to control (Fig. S5C). Interestingly, at 3 h post OMB treatment, the expression status of these 320 proteins was partially reversed, and we observed a significant differential expression of 68 proteins compared to SI animal tissue samples as shown by the Venn diagrams (Fig. 4H and 4I) and heat map analysis (Fig. 4J).

## Discussion

The primary driving force behind systemic oxygenation with OMB treatment is diffusion. Upon delivery to the colon, the OMBs comprising 95-98% oxygen gas deliver oxygen to hypoxic tissue with a diffusivity of ∼2.4 × 10^−6^ cm^2^/s *(13)*. OMBs are superior for delivering oxygen as compared to a macro, non-shelled oxygen gas bubble due to their ability to intimately spread and mix throughout the colon submucosa and increase the interfacial gas-liquid surface area. This enhancement in transport for microbubbles has led to their utilization in industrial fermentation *(14–16)*, and other applications requiring rapid gas absorption. The colon walls are highly vascularized and absorptive (removing ∼ 2 L/day of water from chyme and stool *(17)*) and thereby allow oxygen and carbon dioxide to diffuse across the submucosa from the lumen to adjacent tissue. Within the colon tissue, oxygen diffuses into capillary vessels, binds to deoxygenated blood, and circulates resulting in augmentation of systemic oxygen levels. The statistically significant rises in oxygen blood gas sampled at the carotid and pulmonary arteries demonstrate that OMBs ability can deliver systemic oxygen to a patient via the splanchnic circuit.

The diffusion mechanism responsible for delivering systemic oxygen to an OMB-treated patient is also responsible for removing systemic CO_2_. The permeability of CO_2_ in tissue is ∼20 fold higher than for oxygen *(18)*. Thus, CO_2_ counter-diffuses into the OMB microfoam as O_2_ diffuses into tissue, and the OMB structure is stabilized during this gas-exchange process by the structural integrity of the lipid shell *(19)*. As hypercapnic venous blood passes through the splanchnic circuit, it sheds CO_2_ into the OMB matrix. This lowering of CO_2_ blood gas in the mixed venous return, verified by both P_mv_CO_2_ and ETCO_2_ (Fig. 3B,F,H), reduces the alveolar CO_2_ that would otherwise dilute alveolar O_2_, thereby increasing P_A_O_2_ (Fig. S3). Conversely, all NT animals experienced an increase in CO_2_ levels throughout the study.

In this study, we showed that colonic OMB can improve systemic O_2_ delivery and CO_2_ removal for up to 150 min. This time duration is relevant for acute, severe hypoxia as a possible bridge to alternate therapy, where resources/access to ECMO is limited, or when the risks of pulmonary bypass via established methods outweigh the benefit. Eventually the OMB bolus will deplete its oxygen supply and saturate itself with CO_2_. In such cases, multiple bolus administrations can be administered, where the expired bolus can be flushed naturally or with the help of conventional pro-motility enema. This capability favors colonic administration as the preferred enteral route for OMB therapy. In prior work, we focused on the intraperitoneal (IP) route in small animal lung injury models *(9–11)*. We also examined the IP route here in our SI porcine model and found similar results to the colon route (Figs. S2, Table S2). IP-OMB significantly elevated arterial and mixed venous oxygen tension and simultaneously reduced carbon dioxide over 120 min. IP-OMB increased systemic oxygenation over the NT group (Δ P_a_O_2_ = 17.2 ± 13.9 mmHg within 30 min and Δ P_a_O_2_ = 11.2 ± 4.6 mmHg at 120 min). IP-OMB also decreased systemic CO_2_ levels compared to the NT group (Δ P_a_CO_2_ = -6.4 ± 10.8 mmHg at 30 min and Δ P_a_CO_2_ = -7.9 ± 7.8 mmHg at 120 min). It is plausible that the peritoneal cavity may serve as an additional route to the colon to increase oxygenation further beyond what can be obtained by the colon alone. However, the colon route remains favorable because, for the same OMB dose range, the increase in systemic O_2_ and decrease in CO_2_ was similar for both routes, and the colon route has additional advantages of not requiring surgery to establish a port, the colon can be easily flushed naturally or by enema and re-administered, and it requires only food-grade sterility since it is contained within the gastrointestinal tract.

The main limit of this study is that we examined effects out to a pre-determined 150 min endpoint. Severe hypoxia from respiratory distress may occur for a much longer period, over days and even weeks. Therefore, future work will focus on understanding not only the therapeutic duration of a single OMB bolus, but also the therapeutic effects of multiple doses. Additionally, colonic OMB therapy should be tested in alternative large-animal ARDS models, such as lipopolysaccharide and oleic acid, with varying degrees of severity of lung insult and hypoxia, as well as the interaction between OMB therapy and mechanical ventilation. Colonic OMB therapy should also be investigated as a bridge to ECMO for rapidly deteriorating patients who require additional time for transport and to setup the circuit, or for austere environments with limited resources.

## Materials and Methods

### Animal Subjects

All animal experiments were approved by the University of Nebraska Lincoln (UNL) Institutional Animal Care and Use Committee (IACUC). Female pigs (31-51 kg, n=6 treatment, n=6 no treatment) were housed and cared for according to USDA (United States Department of Agriculture) guidelines. Animals were acclimated to the facility for 4-7 days and received food reward training to ease handling and blood draws. Study animals were fasted overnight and given free access to water for procedures on the following day.

### Animal Preparation and Surgical Procedures

Sedation for peripheral intravenous catheter (PIV) placement and endotracheal intubation was achieved with a mixture of telazol (4.4 mg/kg), ketamine (2.2 mg/kg) and xylazine (2.2 mg/kg) delivered via intramuscular injection. To assist with intubation, an intravenous bolus dose of propofol (2-4.4 mg/kg) was given as needed. Baseline chest x-rays (CXR) (Figs. 1A & 1B) were taken (Portable x-ray unit EPX-F2800, Ecotron Co. Ltc; wireless digital flat panel detector Mars1417V-TSI, iRay Technology, Shanghai, China) prior to smoke inhalation, and at 24 and 48 h after smoke inhalation (Figs. 1D & 1E, at 48 h). An endotracheal tube (#7-8 cuffed; MWI Animal Health, Boise, ID, USA) was inserted into the trachea and animals were ventilated at a tidal volume (TV) of 6-8 mL/kg and peak end expiratory pressure (PEEP) of 5 cmH_2_O (Newport HT70, Medtronic, Minneapolis, MN). Respiratory rate (RR) was adjusted to maintain eucapnia as monitored by end-tidal CO_2_ (ETCO_2_). The fraction of inspired oxygen (FiO_2_) was set at 50% during surgical procedures (central venous catheter placement & arterial catheter placement), then titrated down to 21% and maintained throughout the experiment. Non-invasive monitoring included blood pressure taken by cuff placed around the animal’s hind leg, peripheral oxygen saturation (SpO_2_), heart rate (HR) and ETCO_2_ recorded via the Surgivet monitor (Smiths Medical, Dublin, OH). Continuous IV sedation consisting of midazolam (0.4-0.7 mg/kg/hr), fentanyl (0.03-0.1 mg/kg/hr) and/or propofol (0.2-0.4 mg/kg/min), and maintenance IV fluids (10 mL/kg/hr normal saline) were given throughout the procedure via a quadruple-lumen central venous catheter (8.5 Fr x 16 cm, Arrow International) placed in the internal jugular vein. Core temperature was monitored by rectal probe and a circulating warming blanket was used to prevent body cooling. A urinary catheter was placed to monitor output. Using sterile technique and ultrasound guidance (Butterfly iQ, Butterfly Network, New York City, NY), carotid artery (CA) and femoral artery (FA) access catheters were placed for serial lab draws and invasive blood pressure monitoring (18 GA 16 cm; Femoral Arterial Line Catherization Kit; Teleflex, Morrisville, NC). Pulmonary artery (PA) catheter (8 Fr x 110 cm Swan-Ganz CCOmbo Thermodilution Catheter; Edwards Lifesciences, Irvine, CA) was placed in the internal jugular vein under ultrasound guidance and advanced to the pulmonary artery as confirmed by waveform. The CA and PA access ports were connected to Surgivet monitor and Vigilance II monitor, respectively (Edwards Lifesciences, Irvine, CA) with transducers (Meritans DTXPlus, Disposable Pressure Transducer with EasyVent; Merit Medical, South Jordan, UT, USA). Invasive arterial blood pressure (ABP), central venous pressure (CVP), pulmonary artery pressure (PAP), cardiac output (CO), mixed venous oxygen saturation (S_mv_O_2_), and central (core) temperature were monitored throughout the study. Blood samples were drawn from the CA, FA and PA catheters for the measurement of baseline blood gas prior to smoke inhalation and at pre-determined time intervals throughout the study period (ABL80 FLEX CO-OX, Radiometer, Brea, CA). To maintain patency, catheters were flushed throughout the experiment with 3-5 mL of sterile saline, and a heparin solution (1:500 dilution in 50% dextrose solution) was infused to fill the volume of the port chosen as a “lock” solution. Sedated/anesthetized animals from survival surgeries were continuously monitored until they regained sternal recumbency. All catheters were removed after smoke inhalation was completed. The surgical procedures were repeated at 48 hr after smoke inhalation prior to treatment with OMBs.

### Smoke Inhalation

Upon completion of all surgical procedures, animals (n=12) were exposed to oak wood smoke from a custom-made smoke generator connected in parallel to the endotracheal tube (Fig S1A). The duration of the smoke exposure was 2 hr, and the volume of smoke was approximately 1000 L (as estimated from TV, RR, and total time of smoke exposure). Invasive and noninvasive vital signs were monitored continuously during the experiment. Following smoke exposure, blood samples were collected from CA, FA and PA ports for blood gas analysis. Smoke exposure was stopped immediately if the animal developed hemodynamic instability, which was determined by hypotension (systolic blood pressure less than 60) and irreversible desaturation (SpO_2_ less than 70% despite rescue maneuvers such as increasing FiO_2_).

### Oxygen Microbubble Preparation

OMBs were generated via sonication as described by Feshitan et al.^8^ The resulting OMB solution, which contained 15% oxygen gas by total volume (void fraction (VF) = 0.15), was then centrifuged (Sorvall Legend T, Thermo Scientific, Waltham, MA) as described by Swanson et al.^12^ in batches of four 140 mL syringes (Covidien Monoject 140, Medtronic, Minneapolis, MN) at 300 relative centrifugal force (RCF) for 1 min to achieve a final oxygen gas content of VF ≥ 0.7 (+70% oxygen gas by total volume). The resulting high-concentration oxygen microbubble foam was collected in 2 L gas-tight syringes (S2000, Hamilton, Reno, NV) and stored at ∼ 4 °C. OMB size (Figure 2A) was measured using the electrozone sensing method (Coulter Multisizer III, Beckman Coulter, Opa Locka, FL). OMB VF was calculated by subtracting the weight of a fixed volume of OMB from the weight of an equivalent volume of aqueous solution (lipid-PBS mixture) and dividing by the weight of the aqueous solution at the same fixed volume. Finally, oxygen gas content by total gas volume (%) was measured with an oxygen needle sensor (OX-NP, Unisense, Aarhus, Denmark).

### No Treatment and Colonic Oxygen Microbubble Treatment

At 48 h after smoke inhalation, CXR was obtained and CA, FA, and PA catheters were again placed for serial blood sampling and monitoring as described above. Lung injury from smoke inhalation was confirmed by presence of bilateral diffuse infiltrates on CXR (Figure 1D & E). After completion of catheter placement, FiO_2_ was lowered to 21% and maintained throughout the remainder of the experiment. Other ventilator parameters were set to TV = 6-8 mL/kg, PEEP = 0-1 to maintain normal driving pressure, and RR was adjusted to maintain eucapnia. Baseline blood gas samples (CA, PA and FA ports) were taken every 15 min until the desired level of hypoxia was achieved (P_a_O_2_ ≤ 45 ± 5 mmHg). After three consecutive hypoxic blood gas measurements (P_a_O_2_ ≤ 45 ± 5 mmHg) at 5-min intervals, animals receiving OMB treatment (n=6) had a rectal tube (super xl enema kit, Bracco Diagnostics Inc. Monroe Township, NJ) placed and secured with a purse string suture to ensure a tight seal of the enema tube with the anal canal (Figure 2D). The enema tubing was then connected to the 2L super syringe containing the OMB. The OMB were delivered to the animal at a rate of 500 mL/min which was controlled by a custom syringe pump (Respirogen, Boulder, CO). Three boluses of OMBs were sequentially administered to achieve a 75 – 100 mL/kg total dose (for example, at 100 mL/kg a 45 kg animal would receive a 4.5 L dose). Animals receiving no treatment (n=6) had a theoretical treatment time (t = 0 min) ∼5 min after their last baseline measurement. Following treatment (t = 0 min), arterial and mixed-venous blood gas samples were taken at t = 15, 30, 45, 60, 90, 120, and 150 min post-treatment time. After the experiments, animals were euthanized with an intravenous injection of 0.1 ml/lb of Fatal-Plus (Vortech Pharmaceuticals, Dearborn, MI).

### Plasma Sample Extraction

Blood samples were collected from the CA catheter at baseline, 2h, and 48h time points in lithium heparin BD Microtainer tubes (Becton, Dickinson and Company, Franklin Lakes, NJ). Tubes were immediately inverted 8-10 times to assure anticoagulation and centrifuged at 4000 g for 3 min. Supernatants were collected as plasma samples and stored at -20 °C until analysis. IL-6, IL1β and IL-8 immune assays were performed in samples of 12 animals using IL-6 (catalog#P6000B), IL1β/IL-1F2 (catalog#PLB00B) and IL-8/ CXCL8 (catalog#P8000) Quantikine® ELISA kits, (R&D Systems, Inc., Minneapolis, MN) following manufacturer’s protocol.

### Bronchoalveolar Lavage (BAL)

BAL of pig lungs was performed at baseline, 2h, and 48h time points using a bronchoscope in a set of 12 intubated animals. 10 ml of sterile normal saline was instilled to the secondary and tertiary bronchi through the bronchoscope and ∼5ml of the fluid was collected for analysis. BAL fluid samples were centrifuged immediately at 400 g at 4°C for 10 min and supernatants were at stored at -20 °C until analysis. Total protein quantification was performed in samples using Pierce™ BCA (Bicinchoninic Acid) Protein Assay Kit (Thermo Fisher Scientific Inc. Waltham, MA) following manufacturer’s protocol. IL-6, IL1β and IL-8 immune assays were performed in samples of 12 animals using IL-6 (catalog#P6000B), IL1β/IL-1F2 (catalog#PLB00B) and IL-8/ CXCL8 (catalog#P8000) Quantikine® ELISA kits, (R&D Systems, Inc., Minneapolis, MN) following manufacturer’s protocol.

### Tissue Collection

Necropsy was performed in 15 animals. At necropsy, lung tissues were collected from all five lobes; upper, middle and lower lobes of right lung and upper and lower lobes of left lung for histological examination and pulmonary edema assessment. Tissues for histology were immediately placed in 10% neutral buffer formalin fixative for approximately 24 h. Formalin fixed tissues were placed into 70% ethanol and transferred to the University of Nebraska Medical Center (UNMC) Tissue Science Facility (TSF) for further tissue processing and embedment in paraffin blocks.

### Lung Injury Score

The lung tissue (n=15) in 10% neutral formalin was dehydrated in graded concentrations of ethanol solution and cleared in xylene. The tissue samples were then paraffin-embedded, sectioned with 4-μm thickness, and stained with hematoxylin and eosin at the UNMC Tissue Sciences Facility using automated Ventana Discovery Ultra (Roche Diagnostics, Indianapolis, IN) as per manufacturer’s protocol. An independent pathologist performed a blinded examination of the tissues under light microscope. Ten fields of each lung tissue section were examined at magnification x400. The severity of the lung injury was scored by the criteria of alveolar edema, intra-alveolar hemorrhage, and leukocyte infiltration. Alveolar edema and intra-alveolar hemorrhage were scored on a scale from 0 to 3; where 0 = < 5% of maximum pathology, 1 = mild (< 10%), 2 = moderate (10 - 20%), and 3 = severe (20 - 30%). Leukocyte infiltration was also scored on a scale from 0 to 3; where 0 = absence of extravascular leukocytes, 1 = < 10, 2 = 10 - 45, and 3 = > 45 leukocytes.

### Lung Tissue Lysate Preparation

Fresh frozen lung lobe tissues with highest injury score (n=9) were homogenized using VWR® Mini Bead Mill Homogenizer (VWR International LLC., Radnor, PA) following manufacturer’s protocol. Briefly, frozen tissues of three control, three SI animals and three SI+OMB animals were washed in cold X1 PBS, and 30 mg of each tissue was placed separately in a 2 mL tube containing 2.8 mm ceramic beads and 750 μl of lysis buffer containing RIPA buffer (Thermo Fisher Scientific Inc. Waltham, MA) and protease inhibitor cocktail (Sigma Aldrich Inc., St. Louis, MO) at room temperature. The samples were homogenized at speed 4 for 60 seconds. This was followed by incubation in ice for 30 min and centrifugation at 13,000 rpm for 20 min at 4 °C. Protein concentration was determined using Pierce™ BCA (Bicinchoninic Acid) Protein Assay Kit (Thermo Fisher Scientific Inc. Waltham, MA) following manufacturer’s protocol.

### Immunoblot Analysis

Protein (50 μg) was separated by SDS□ polyacrylamide gel electrophoresis and transferred onto PVDF (polyvinylidene fluoride) membrane (Bio-Rad Lab Inc., Hercules, CA) by electro blotting. The PVDF membrane was blocked with 5% nonfat dry milk in X1 TBST (50mM Tris, pH 7.5, 150mM NaCl, 0.01% Tween 20) for 1 h at room temperature (RT). The membrane was then incubated in primary antibody, IL-6 antibody (#ab6672, Abcam Inc, Cambridge, MA) and IL-1β (#P420B, Thermo Fisher Scientific, Waltham, MA) or β-actin (#4970, Cell Signaling Technology Inc., Danvers, MA) at 1:1000 dilution in X1 TBST with 5% bovine serum albumin (Sigma Aldrich Inc., St. Louis, MO) overnight at 4°C. The membrane was washed three times with X1 TBST for 10 min each and incubated with HRP□conjugated rabbit or mouse secondary antibodies (#7074 and #7076, Cell Signaling Technology Inc., Danvers, MA) at 1:5000 dilution in X1 TBST with 5% nonfat dry milk for 1 h at RT. Following three washes in X1 TBST, proteins were detected using the enhanced chemiluminescence system (Bio-Rad Lab Inc, Hercules, CA) and image with ChemiDoc™ MP Imaging System (Bio-Rad Lab Inc, Hercules, CA)

### Sample preparation for mass spectrometry

The protein concentration in the cell lysates was estimated using BCA Protein Assay Kit (Pierce) for each sample. The protein digestion for mass spectrometry and TMT labelling of the peptides were carried out following the manufacturer’s suggestions. Briefly, 100 μg of proteins from each lysate was reconstituted to 100 μL with 100 mM triethylammonium bicarbonate (TEAB). Proteins were next reduced with 5 μL of 200 mM tris(2-carboxyethyl) phosphine (TCEP) (1 h incubation, 55 □C) and alkylated with 5 μL of 375 mM iodoacetamide (IAA) (30 min incubation in the dark, room temperature). The reduced and alkylated proteins were purified with acetone precipitation at -20 □C overnight. The protein precipitates were collected by centrifugation at 8000 × g for 10 min at 4 □C. The pellets were air-dried and resuspended in 100 μL of 50 mM TEAB. Next, the protein digestion was carried out using 2.5 μg of trypsin per sample (24 h incubation, 37 □C). The amount of peptide yielded in each sample was estimated with the Pierce Colorimetric Peptide Assay kit. The amounts of peptides to be tagged were normalized and mixed with 41 μL of TMT reagent (TMT sixplex, Thermo Fisher Sci) freshly dissolved in acetonitrile (20 μg/μL) (1 h incubation, room temperature). 8 μL of 5% hydroxylamine was added to quench the reaction (15 min incubation, room temperature). Tagged tryptic peptides were pooled and concentrated to around 20 μL by vacuum centrifugation and analyzed using a high-resolution mass spectrometry nano-LC-MS/MS Tribrid system, Orbitrap Fusion™ Lumos™ coupled with UltiMate 3000 HPLC system (Thermo Scientific).

### LC-MS/MS and bioinformatics analysis

Around 1 µg of peptides were run on the pre-column (Acclaim PepMap™ 100, 75μm × 2cm, nanoViper, Thermo Scientific) and the analytical column (Acclaim PepMap™ RSCL, 75 μm × 50 cm, nanoViper, Thermo Scientific). The peptides were eluted using a 125-min linear gradient of ACN (0-45 %) in 0.1% FA and introduced to the mass spectrometer with a nanospray source. The MS scan was done using detector: Orbitrap resolution 120000; scan range 375-1500 m/z; RF lens 60%; AGC target 5.0e5; maximum injection time 150 ms. Ions with intensity higher than 5.0e3 and charge state 2-7 were selected in the MS scan for further fragmentation. MS2 scan parameters set: CID collision energy 35%; activation Q 0.25; AGC target 1.0e4; maximum injection time 150 ms. MS3 scan parameters were set: HCD collision energy 65%; Orbitrap resolution 50000; scan range 100-500 m/z; AGC target 1.0e5, maximum injection time 200 ms. All MS and MSn collected spectra were analyzed using Protein Discoverer (Thermo Fisher Sci, vs 2.2.) pipeline. Sequest HT was set up to search the NCBI database (selected for Sus scrofa, 2019_01, 63657 entries), assuming the digestion enzyme trypsin. The parameters for Sequest HT were set as follows: Enzyme: trypsin, Max missed cleavage: 2, Precursor mass tolerance: 10 ppm, Peptide tolerance: ± 0.02 Da, Fixed modifications: carbamidomethyl (C), TMT sixplex (any N-terminus); Dynamic modifications: oxidation (M), TMT sixplex (K). The parameters for Reporter ions quantifier were set as follows: Integration tolerance: 20 ppm, integration method: most confident centroid, mass analyzer: FTMS, MS order: MS3, activation type: HCD, min. collision energy: 0. max. collision energy: 1000. Percolator was used to calculate the false discovery rate (FDR) for the peptide spectral matches. The parameters for Percolator were set as follows: target FDR (strict): 0.01, target FDR (relaxed): 0.05, validation based on: q-value. Quantification parameters were set: Peptides to use: unique+razor, Normalization mode: total peptide amount.

### Statistics Analysis

All oxygen and CO2 blood gas data are reported as mean ± standard deviation. Statistical significance was based on an unpaired parametric t-test with Welch correction (non-equal standard deviations) between the delta of the no treatment group (n=6) and the delta of the OMB treatment group at the 15, 30, 45, 60, 90, 120 and 150 min time points (Table S1) and out to 120 min for the intraperitoneal OMB treatment group (Table S2). The colonic OMB treatment group fell to an n=5 after 60 min due to a loss in data collection for one of the animals. The same animal had a loss in collection of PaCO2 data within 15 min of OMB treatment. An additional animal dataset did not contain PmvCO2 data due to equipment error bringing the PmvCO2 sample size from n=5 to n=4 after 60 min. The intraperitoneal OMB treatment group had a n=5 due to the study switching to the colonic route of administration after the 5^th^ animal. One-way ANOVA for multiple comparisons was used to compare control (n=4) vs. SI (n=5) vs. SI+OMB (n=5) samples for lung injury and wet/dry ratios. IL-1ß immunoblot analysis was performed on n=3 baseline samples and n=4 NT and OMB samples. IL-6 marker injury analysis was performed on n=11 animals for both baseline and 2-hour post-smoke inhalation time points for both BAL and Plasma samples. The IL-6 treatment analysis was performed on n=6 BAL samples and n=5 Plasma samples for both the NT and OMB treatment groups 48-hours post-SI injury.

## Supplementary Materials

### Materials and Methods

Fig. S1. Smoke Inhalation Injury and Intraperitoneal OMB Delivery Schematics

Fig. S2. Intraperitoneal Oxygen and CO2 Blood Gas Measurements

Fig. S3. Alveolar Oxygen Pressure

Fig. S4. IL-6 Immunoblot and IL-8 and IL-1ß ELISA Results

Fig. S5. Differential Global Protein Expressions

Table S1. Animal Oxygen Measurements and Counts

Table S2. Title of the first supplementary table

Movie S1. OMB Colonic Administration

## Acknowledgments

We would also like to acknowledge Andrew Kingsbury for generating the schematics presented in this manuscript.

## Author contributions

Conceptualization: PAM, RMS, MAB, KLB

Methodology: PAM, PDL, RTS, MAB, KLB

Investigation: PAM, PDL, HRW, AM, RTS, EMD, JC, KLB

Visualization: PAM, PDL, HRW, MAB, KLB

Funding acquisition: MAB, KLB

Project administration: PAM, RMS, MAB, KLB

Supervision: PAM, MAB, KLB

Writing – original draft: PAM, PDL

Writing – review & editing: PAM, PDL, MAB, KLB

