## Supplementary figures and images for "Colonic Oxygen Microbubbles Augment Systemic Oxygenation and CO_2_ Removal in a Porcine Smoke Inhalation Model of Severe Hypoxia"

### Fig. S1.

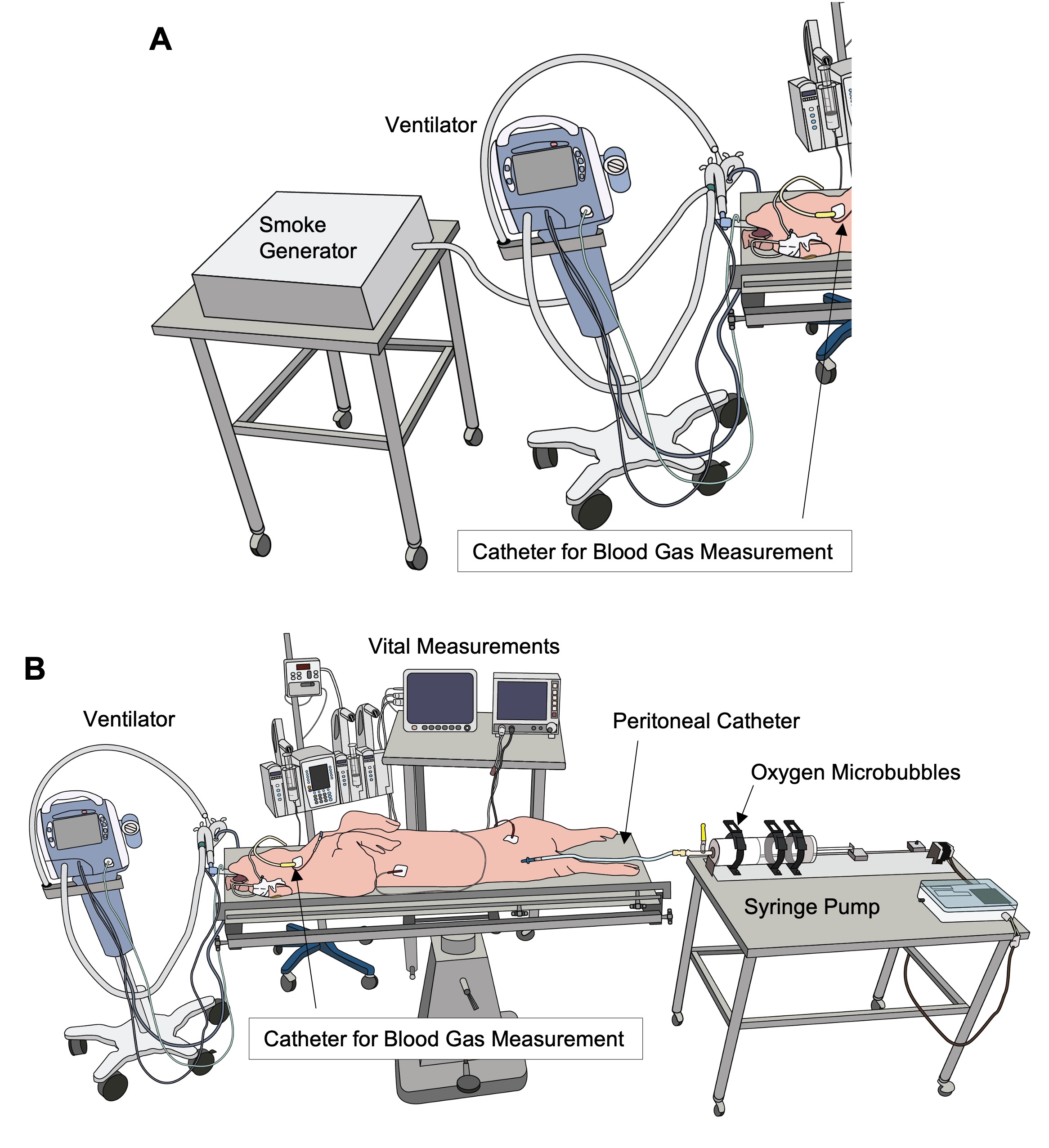

### Fig. S2.

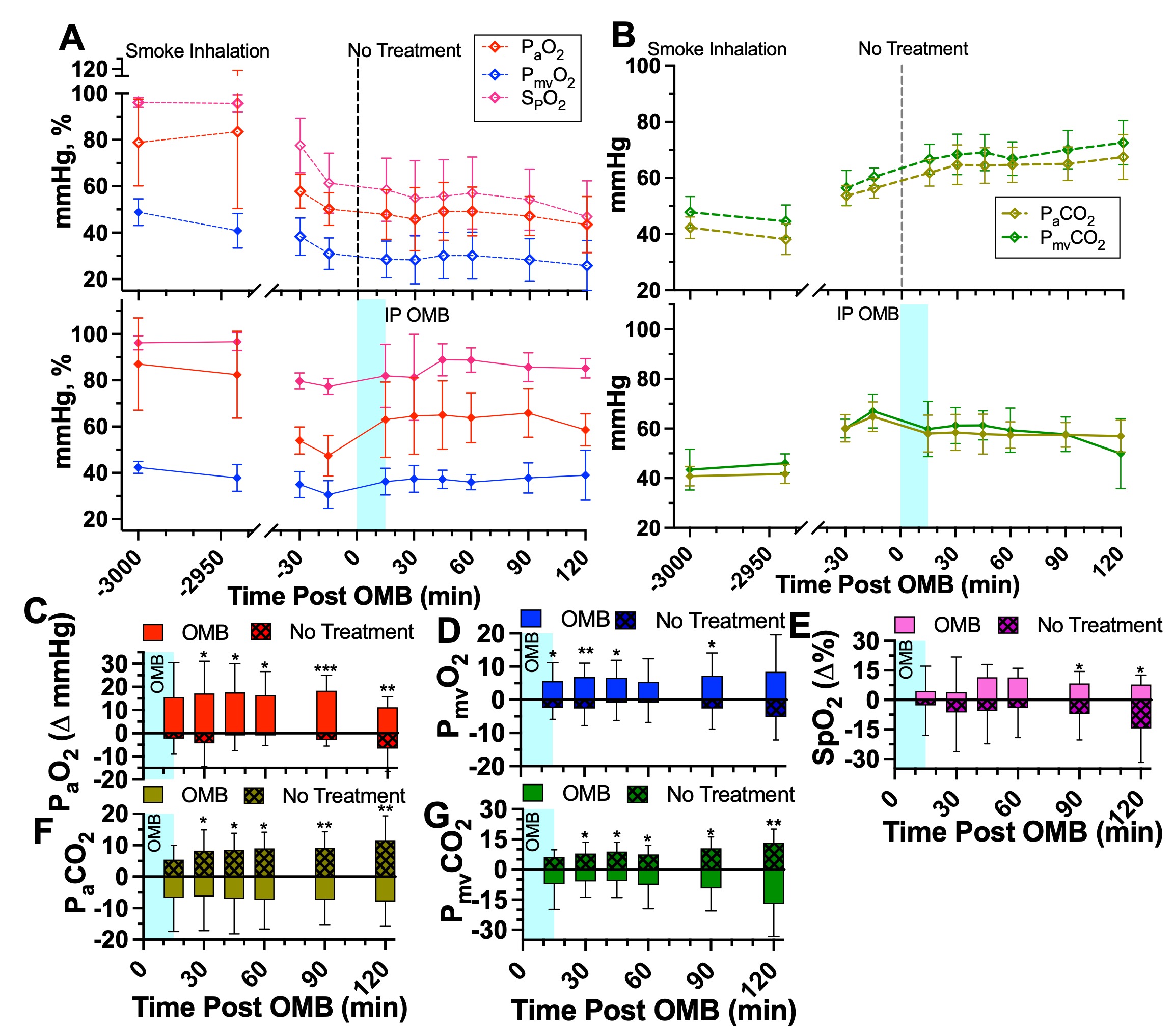

### Fig. S3

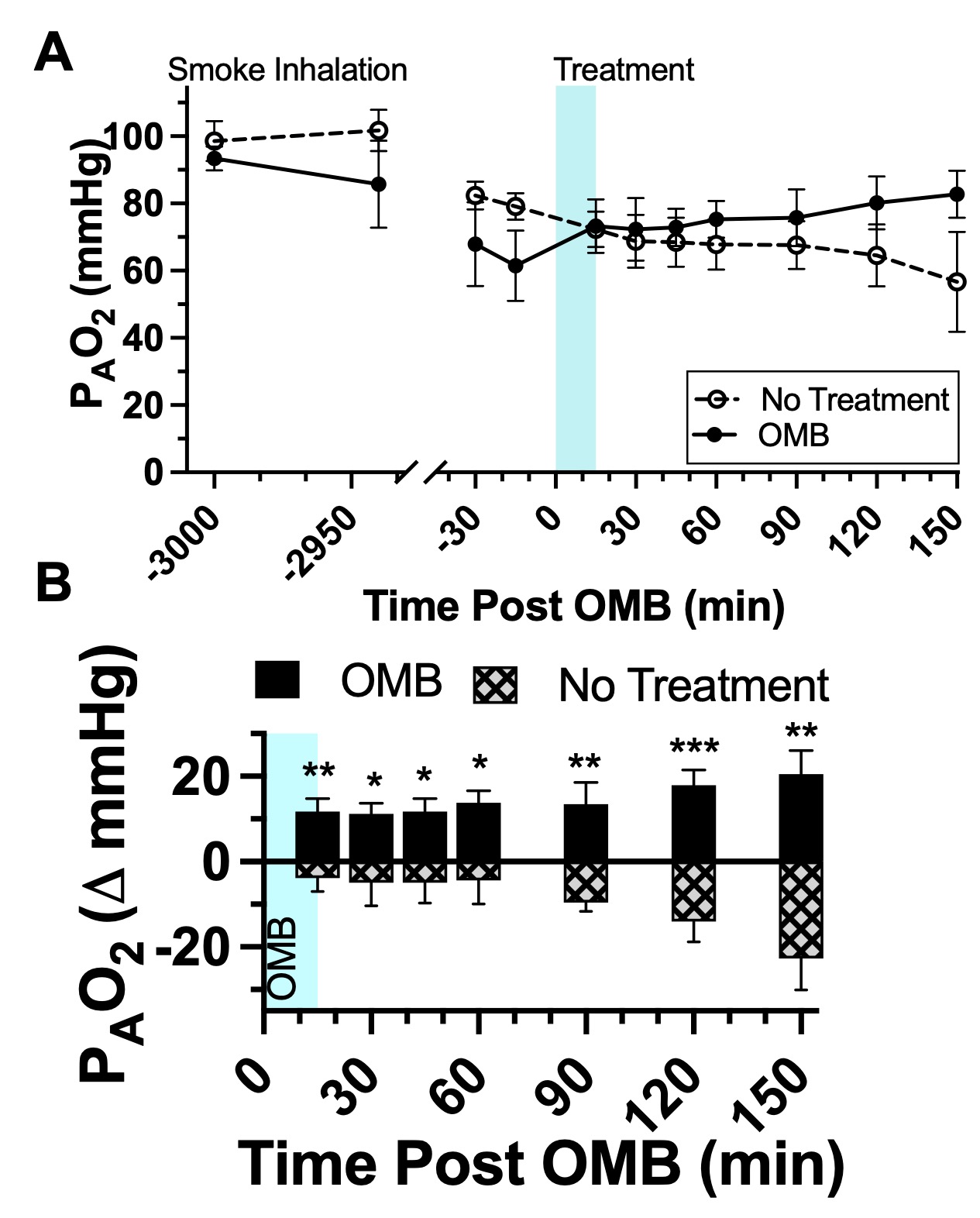

### Fig. S4.

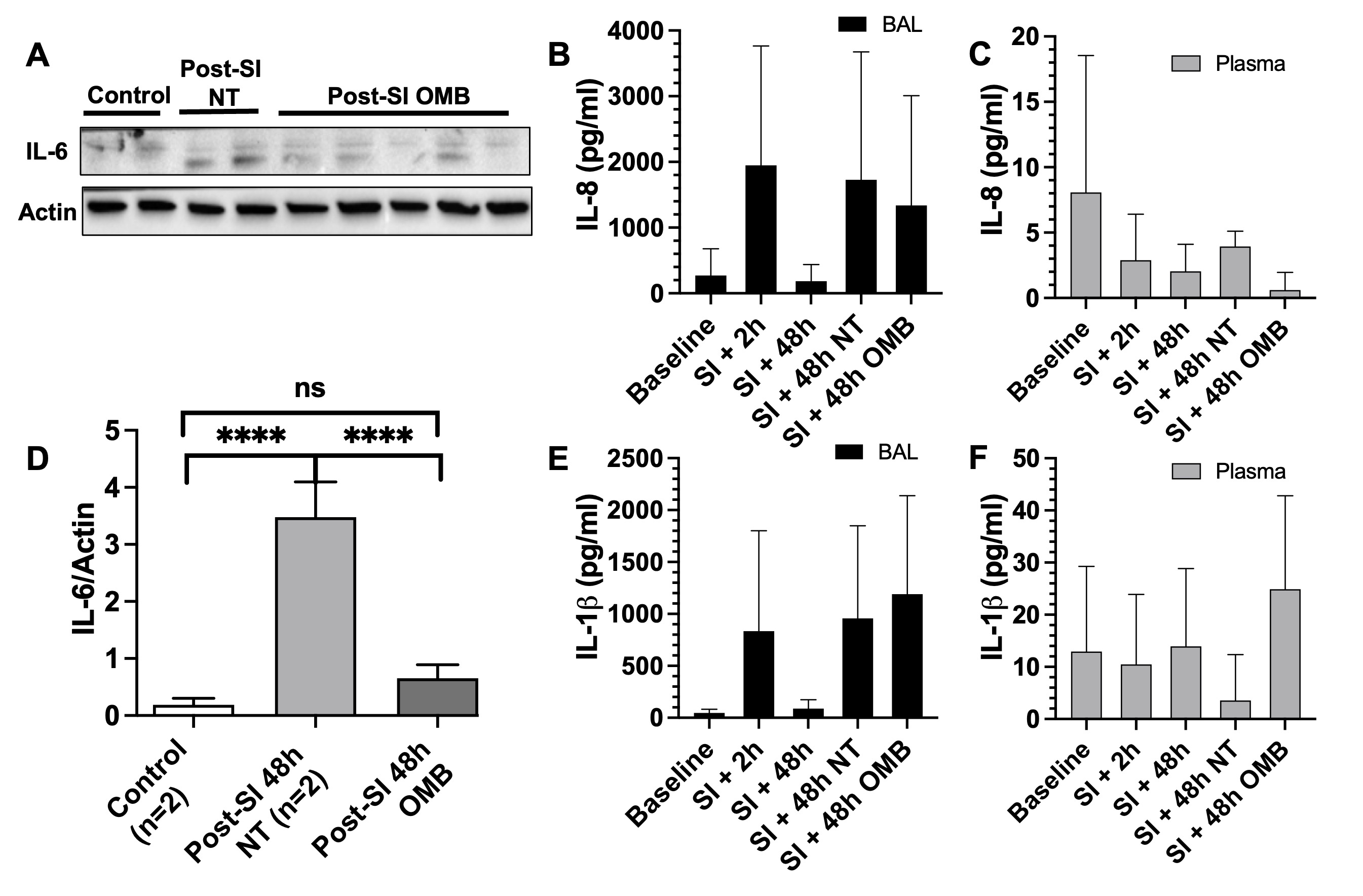

### Fig. S5.

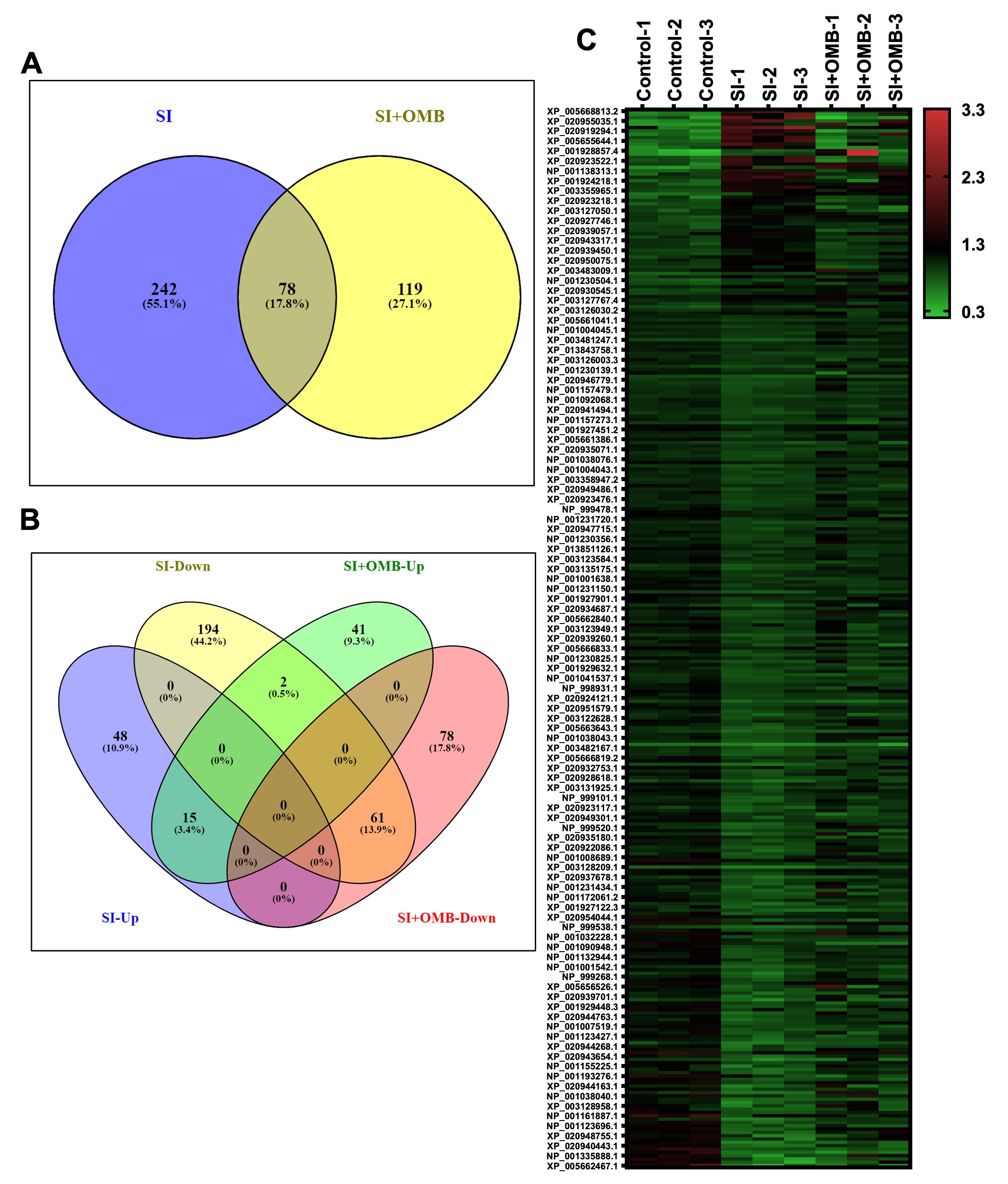

### Table S1.

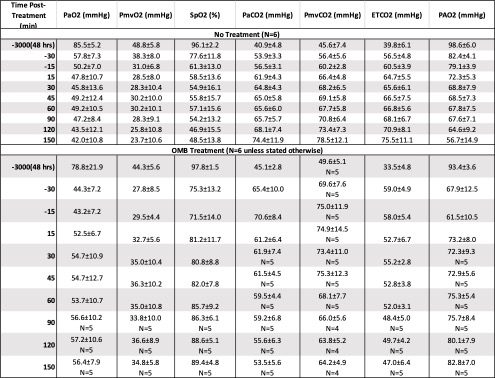

### Table S2.

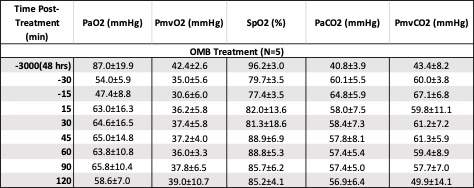
