## Supplemental Materials and Methods for "Colonic Oxygen Microbubbles Augment Systemic Oxygenation and CO_2_ Removal in a Porcine Smoke Inhalation Model of Severe Hypoxia"

*Intraperitoneal Oxygen Microbubble Treatment*

After the development of ARDS 48 h post SI, animals were placed under general anesthesia as before. Animals receiving a peritoneal dose of OMB had an intraperitoneal catheter placed. Specifically, IP catheters were placed in the lower right-hand quadrant of the abdomen while retracting the skin over the insertion point, using a Chest Tube with Sharp Trocar (MILA International, #CTT28, 28Fr x 42cm) to create an opening in the abdominal wall. The skin was first sanitized with chlorhexidine (2%) and isopropyl alcohol (70%). SpO2 and vital signs were monitored, and animals were maintained on minimal ventilator settings with FiO2 set at 21%. After three consecutive hypoxic blood gas measurements (PaO2≤45 +/- 5 mmHg) at 5-min intervals, animals were given a one-time bolus treatment of OMB via the peritoneal catheter (dose volume of 3.6-4.3 L) (Fig. S1B). Following treatment (t = 0 min), arterial and mixed-venous blood gas samples were taken at t = 15, 30, 45, 60, 90, and 120 min post-treatment time. Animals were euthanized and lung tissue collected for analysis at the end of the study.

*Alveolar Oxygen Pressure Quantification*

The alveolar pressure of oxygen in the lungs (P_A_O_2_) is the estimated amount of oxygen in the alveoli. The P_A_O_2_ for study animals was calculated with the equation^13,14^:

$P_{A}O_{2}=\left( P_{b}+MAP-P_{H_{2}O} \right)*FiO_{2}-\frac{P_{a}CO_{2}}{\mathrm{RQ}}$	[1]

where P_b_ is the atmospheric pressure (760 mmHg), MAP is the mean airway pressure, P_H2O_ is the water vapor pressure (47 mmHg) and RQ is the respiratory quotient (0.8).
