## Supplemental Figures and Tables for "Colonic Oxygen Microbubbles Augment Systemic Oxygenation and CO_2_ Removal in a Porcine Smoke Inhalation Model of Severe Hypoxia"

**Fig. S1. Smoke Inhalation Injury and Peritoneal OMB Delivery. A)** Smoke inhalation injury diagram depicting the smoke generator providing the ventilator with smoke and delivering to the animal via the lungs. **B**) Diagram showing the delivery of OMB to a smoke inhalation lung injury pig via an intraperitoneal catheter to the peritoneal cavity.

**Fig. S2. Intraperitoneal OMB Oxygenation and CO2 Removal. A)** Blood oxygen and **B**) CO_2_ measurements for NT (top) and intraperitoneal OMB (bottom) treatment groups from before smoke inhalation injury (t = -3000 min) out to t = 150 min post treatment time. The change in oxygen vitals for NT (hashed) and OMB (solid) treatment groups showing the statistical significance at times t = 15, 30, 45, 60, 90 and 120 min ((**C**) P_a_O_2_ (red), (p=0.0515, 0.0226, 0.0243, 0.0155, 0.0010 and 0.0049), (**D**) P_mv_O_2_ (blue), (p=0.0280, 0.0084, 0.0480, 0.1550, 0.0376 and 0.0535) (**E**) SpO_2_ (violet), (p=0.4017, 0.3897, 0.0566, 0.0502, 0.0365 and 0.0243), (**F**) P_a_CO_2_ (gold), (p=0.0606, 0.0364, 0.0323, 0.0125, 0.0055 and 0.0027), and (**G**) P_mv_CO_2_ (green), (p=0.0714, 0.0138, 0.0117, 0.0406, 0.0120 and 0.0098).

**Fig. S3. Colonic OMB Alveolar Oxygen Pressure. A**) P_A_O_2_ measurements for NT (dashed) and colonic OMB (solid) treatment groups from before smoke inhalation injury (t = -3000 min) out to t = 150 min post treatment time. The change in P_A_O_2_ for NT (hashed) and OMB (solid) treatment groups showing the statistical significance at times t = 15, 30, 45, 60, 90, 120 and 150 min (p=0.0046, 0.0328, 0.0186, 0.0201, 0.0078, 0.0009 and 0.0018).

**Fig. S4. IL-6 Immunoblot and IL-8 and IL-1β ELISA analysis. A** and **D)** Immunoblot analysis of IL-6 expression levels in fresh frozen lung tissues of Control (t = -48 h), SI (t = -46 h) and SI+OMB (t = 3 h) groups (p<0.0001). **B** and **C**) IL-8 expression level in BAL fluid and plasma samples of treated animals at baseline (t = -48 h), SI+2h (t = -46 h), SI+48h (t = -0.5 h), NT and OMB (t = 3 h) samples. **E** and **F**), IL-1β expression level in BAL fluid and plasma samples of treated animals at the same timepoints as IL-8 analysis.

**Fig. S5. Additional Proteomics Data.** Venn diagram showed 320 proteins with significant differential expression between SI and control groups (**A** and **B**). **C**) Heat map analysis of 320 proteins that were significantly upregulated or downregulated between SI and control groups and their corresponding expression at 3 hours post OMB treatment.

**Table S1. NT and Colonic OMB Sample Sizes and oxygen and CO2 measurements.** Animal oxygen and CO_2_ blood gas measurements and counts for time points t = -3000, -30, -15, 15, 30, 45, 60, 90, 120 and 150 min for NT and colonic OMB treatment groups.

**Table S2. Peritoneal OMB Sample Sizes and Oxygen and CO2 Measurements.** Animal oxygen and CO_2_ blood gas measurements and counts for time points t = -3000, -30, -15, 15, 30, 45, 60, 90 and 120 min for the intraperitoneal OMB treatment group.
